# *FLOWERING LOCUS T4 (HvFT4)* delays flowering and decreases floret fertility in barley

**DOI:** 10.1101/2020.03.26.010033

**Authors:** Rebecca Pieper, Filipa Tomé, Maria von Korff

**Affiliations:** Institute for Plant Genetics, Heinrich-Heine University Düsseldorf, Düsseldorf, Germany; Max-Planck-Institute for Plant Breeding Research, Cologne, Germany; Cluster of Excellence on Plant Sciences, “SMART Plants for Tomorrow’s Needs”, Heinrich-Heine University Düsseldorf, Düsseldorf, Germany

**Keywords:** flowering, fertility, FLOWERING LOCUS T, barley, photoperiod, cereals

## Abstract

*FLOWERING LOCUS T*-like genes (*FT*-like) control the photoperiodic regulation of flowering in many angiosperm plants. The family of *FT*-like genes is characterised by extensive gene duplication and subsequent diversification of *FT* functions which occurred independently in modern angiosperm lineages. In barley, there are 12 known *FT*-like genes (*HvFT*) but the function of most of them remains uncharacterised. This study aimed to characterise the role of HvFT4 in flowering time control and development in barley. The overexpression of *HvFT4* in the spring cultivar Golden Promise delayed flowering time under long-day conditions. Microscopic dissection of the shoot apical meristem (SAM) revealed that overexpression of *HvFT4* specifically delayed spikelet initiation and reduced the number of spikelet primordia and grains per spike. Furthermore, ectopic overexpression of *HvFT4* was associated with floret abortion and with the downregulation of the barley MADS-box genes *VRN-H1, HvBM3* and *HvBM8* which promote floral development. This suggests that HvFT4 functions as a repressor of reproductive development in barley. Unraveling the genetic basis of *FT*-like genes can contribute to the identification of novel breeding targets to modify reproductive development and thereby spike morphology and grain yield.

**Highlight:** We identify the *FLOWERING LOCUS T* (*FT*)-like gene *HvFT4* as a negative regulator of reproductive development, spikelet initiation, floret fertility and grain number in barley.

## Introduction

Variation in flowering time was crucial for the successful adaptation of crop plants to many different geographic areas and strongly impacts yield and reproductive success (Turner et al., 2005; Cockram et al., 2007; Verhoeven et al., 2008; Comadran et al., 2012; Campoli and von Korff, 2014; Gol et al., 2017).

Flowering time is a complex trait regulated by environmental (photoperiod, ambient temperature, vernalization) and internal cues (autonomous, circadian clock, age, gibberellin, sugar availability) (Mouradov et al., 2002; Fornara et al., 2010; Andrés and Coupland, 2012). In Arabidopsis, the different endogenous and environmental cues are integrated by the key floral integrator *FLOWERING LOCUS T (FT)* (Kardailsky et al., 1999; Kobayashi et al., 1999). *FT* is expressed under long day (LD) conditions in the leaves and then translocated as a protein to the shoot apical meristem (SAM), where it interacts with the basic leucine zipper (bZIP) transcription factor *FLOWERING LOCUS D (FD)* to activate the expression of meristem identity genes to induce floral transition (Abe et al., 2005; Wigge et al., 2005; Corbesier et al., 2007). Large *FT*-like gene families have been found in the genomes of cereal monocots such as wheat and barley (12 FT paralogs each), rice (13 FT paralogs) and maize (15 FT paralogs) (Chardon and Damerval 2005, Danilevskaya et al., 2008, Halliwell et al., 2016). Several studies have demonstrated that these *FT*-like gene families arose from gene duplication events followed by subfunctionalisation or neofunctionalization within and between species. One example of subfunctionalisation of FT-like paralogs can be found in perennial poplar trees (*Popolus* spp.), where *FT1* and *FLOWERING LOCUS T2* (*FT2*) have functionally diverged to coordinate flowering and growth-cycles (Hsu et al., 2011). Furthermore, functional diversification has also been demonstrated in rice, where two FT1-like paralogs, *HEADING DATE3a* (*Hd3a*) and *Rice FT1* (*RFT1*), act as photoperiod-specific florigens. *Hd3a* is induced under inductive short day (SD) conditions to promote flowering, in contrast to its closest homolog *RFT1* which acts as major floral activator under LD conditions (Kojima et al., 2002; Hayama et al., 2003, Tamaki et al., 2007; Komiya et al., 2008). Similarly, in barley *FT1* and *FT2* are expressed under LDs and promote floral development, while *FT3* is expressed under SDs and LDs and induces spikelet initiation (Digel et al., 2015; Mulki et al., 2018; Shaw et al., 2019). However, the multifaceted roles of *FT*-like genes after extensive gene duplication events within most flowering species is not yet described.

Barley is a facultative LD plant with either a winter or spring growth habit. Growth habit and vernalization requirement are determined by the genetic interaction of *VERNALIZATION 1* (*VRN-H1*) and *VERNALIZATION 2* (*VRN-H2*). *VRN-H2* encodes a zinc finger and CCT domain (for CONSTANS [CO], CONSTANS-LIKE [CO-like], and TIMING OF CAB EXPRESSION1 [TOC1]) containing protein and is expressed under LDs before winter (Yan et al., 2004). The *APETALA1* (*AP1*)/*CAULIFLOWER* (*CA*L)/*FRUITFULL* (*FUL*)-like MADS box transcription factor *VRN-H1* is a repressor of *VRN-H2* and its upregulation during vernalization releases *HvFT1* and *HvFT3* expression (Yan et al., 2003; 2004; Mulki et al., 2018). Allelic variation in *VRN-H1* (deletions in the regulatory regions of the first intron) and *VRN-H2* (deletion of the gene locus) induce a vernalization-independent expression of *HvFT1* resulting in a spring growth habit (Hemming et al., 2009, Rollins et al., 2013). Photoperiod response is controlled by *PHOTOPERIOD 1* (*PPD-H1*), a homolog of the Arabidopsis PSEUDO RESPONSE REGULATOR proteins with a pseudo-receiver and a CCT domain (Turner et al., 2005). Under inductive day length (LD), *PPD-H1* activates *HvFT1* transcription in the leaves and thereby accelerates flowering (Laurie et al., 1995; Turner et al., 2005; Yan et al., 2006; Hemming et al., 2008). Spring barley varieties carry a mutation in the CCT domain of *Ppd-H1* that is associated with a decreased *HvFT1* expression level under LD conditions and a delay in flowering (Turner et al., 2005; Hemming et al., 2008). While *HvFT1* is only expressed and induces flowering under LD conditions, the homolog *HvFT3* is expressed and promotes reproductive development under long and short-day conditions (Mulki et al., 2018). Experiments in wheat, barley, Brachypodium and rice suggested that *FT2* is a floral promoter, downstream of *FT1* and expressed in the leaf as well as in the inflorescence (Kikuchi et al., 2009; Lv et al., 2014; Digel et al., 2015; Shaw et al., 2019).

The role of the barley *FT* paralogues *HvFT4* to *HvFT12* is yet undescribed and it is not clear if or how these *FT*-like genes differ in function. This study aimed to functionally characterise the role of *HvFT4* in flowering time and reproductive development in barley. Specific goals were to characterise the effects of *HvFT4* overexpression on macroscopic and microscopic inflorescence development under LDs and study the pleiotropic effect of *HvFT4* overexpression on vegetative and reproductive traits. Another objective was to identify target genes of *HvFT4* by analyzing the expression of main flowering time genes in the leaves and inflorescence in response to *HvFT4* overexpression. Finally, the amino acid sequence of HvFT4 was compared to several known floral repressors and promoters from diverse species to identify common amino acid changes in conserved motifs related to flowering time control. This study demonstrates that overexpression of *HvFT4* delayed flowering specifically by delaying spikelet initiation, negatively affected fertility and grain and tiller number.

## Materials and Methods

### Generation of transgenic *Ubi::HvFT4* lines

Transgenic *Ubi::HvFT4* lines were generated as described for *Ubi::HvCO1* (Campoli et al., 2012a), *Ubi::HvCO2* (Mulki et al., 2016) and *Ubi::HvFT3* (Mulki et al., 2018). The *HvFT4* fragment was cloned from cDNA (cv. Optic) and is identical to the Morex sequence DQ411320.

### Plant material and growth conditions

Three independent transgenic T1 and T2 families designated *Ubi::HvFT4* lines 491 (OX-491), 483 (OX-483) and 517 (OX-517) were sown in 96-well growing trays (Einheitserde, 100 mL/cell) together with a null segregant line (null) and the wild-type spring cultivar Golden Promise (WT). The null segregant sister line without the transgene was used as a control together with Golden Promise. The grains were stratified at 4°C for three days for even germination and then transferred to the greenhouse (LD, 16h light/8h darkness) and controlled temperatures (20°C/16°C days/nights). Germination was recorded as the day of coleoptile emergence from the soil and the plants were transferred to single pots (Einheitserde, 1 L/pot) 14 days after emergence (DAE). Repotted plants were randomised following a random block design.

### Genotyping by polymerase chain reactions (PCR)

To confirm the *Ubi::HvFT4* insertion, genomic DNA was extracted following the Biosprint DNA extraction protocol (Qiagen) and was eluted in 200 µl of deionized water. *Ubi::HvFT4* plants were screened for the presence of the transgene using primers that amplify the hygromycin selectable marker gene (Vec8_F/Vec8_R), located on the transformation vector, and the HvFT4 cDNA sequence (FT4_tg_1F/nos_tg_1R), but not the *HvFT4* genomic DNA (Supplementary Table 1). Amplifications were performed in 1x Green GoTaq Reaction Buffer (Promega) with 5 µl of 1:3 diluted template DNA, 0.5 U GoTaq G2 DNA Polymerase (Promega), 64 µM PCR Nucleotide Mix (Promega), 1.5 mM MgCl2 and 0.2 µM of upstream and downstream primer each. Conditions were as follows: 95°C (3 min), 34 cycles of 95°C (30 sec), 57.5°C (1 min), 72°C (1 min) and 72°C (10 min). 12 µl of the PCR product were visualized on a 2 %-agarose gel with ethidium bromide as dye.

### Confirmation of *HvFT4* overexpression in transgenic *Ubi::HvFT4* lines

To confirm *HvFT4* overexpression, the middle part of the youngest fully emerged leaf of main shoots was collected 12 days after emergence (DAE) ten hours after lights-on (zeitgeber time T10). Samples were immediately frozen in liquid nitrogen and stored at −80°C until RNA extraction and real-time quantitative reverse transcription-PCR as described below.

### Phenotyping

Phenotypes were recorded for six to 32 plants of each of the transgenic line, the null segregant and Golden Promise. Flowering time was measured in days from emergence until heading date. Heading was scored as the appearance of 1 cm of the awns from the main shoot flag leaf sheath (Zadoks stage 49) (Zadoks et al., 1974). Morphological phenotypes of the shoot were recorded at heading such as the number of tillers, number of leaves on the main culm, leaf size, and plant height or at plant maturity such as peduncle extrusion. The leaf size (width and length) of the flag leaf and of the three youngest leaves before the flag leaf (Leaf A, B, C) was measured on the main culm. The leaf width was measured at the widest point of the leaf blade and the leaf length was measured from the ligule to the tip. The height of the plants was measured as the distance from the crown to the collar of the main shoot flag leaf. Peduncle extrusion of the main shoot was measured as the distance from the collar of the flag leaf to the base of the spike.

Plants were harvested individually and the total number of florets on the main spike as well as the number of grains on each rachis node of the main spike were recorded. Floret fertility was calculated as the relative number of grains compared to the total number of central florets. In addition, the number of fertile tillers (fertile = at least one grain) was counted. Tiller fertility was calculated as the relative number of fertile tillers compared to the total number of tillers. Total shoot dry mass and fertile spike dry mass was measured to calculate the Harvest Index (HI) as follows, HI = “Fertile spikes dry mass [g]” /”Total shoot dry mass [g]”. The grains from the main shoot were cleaned and grain width, length and area and thousand grain weight (TGW) were measured with the MARVIN Seed Analyser (GTA Sensorik).

### Microscopic inflorescence development and gene expression during plant development

One *Ubi::HvFT4* line (OX-517), the null segregant control, and the wild-type Golden Promise (WT) were cultivated and genotyped as described above. Three primary shoots per genotype were dissected with a microsurgical knife (5 mm blade, Surgical Specialties Corporation) under the stereo microscope every seven days starting at 21 DAE. The development of the meristem was scored according to the quantitative Waddington scale (Waddington et al., 1983). A stereo microscope (Nikon SMZ18) equipped with a digital camera (Nikon digital sight DS-U3) was used to obtain images of apices. For apex sampling, the surrounding leaves were removed from the main shoot apex (MSA) and the apex was cut from the stem with the microsurgical knife. Meristematic tissue was collected at Waddington stages 3.5-4.5 and 6.0-7.0 by pooling 5 meristems and was immediately frozen in liquid nitrogen. In addition, a leaf sample from the selected plants was taken before dissection, as described above, and pooled in the same way as the meristems. All samples were taken at zeitgeber time T10.

### RNA extraction, cDNA synthesis and real-time quantitative reverse transcription-PCR (qRT-PCR)

Leaf and meristem material for gene expression analysis was ground and subjected to RNA extraction using TRIzol reagent (Invitrogen) according to the manufacturer’s instructions with subsequent DNase I treatment (Thermo Scientific). First-strand cDNA synthesis was performed using approximately 4 µg of total RNA with 0.5 mM dNTP Mix (Thermo Scientific), 1 µg Oligo(dT)12-18 primer (metabion international AG), 0.01 M DTT (Invitrogen) and 150 U SuperScript® II Reverse Transcriptase (Invitrogen) in 1x First-Strand Buffer (final Volume 40 µl) following the manufacturer’s instructions and subsequently 1:4 diluted in nuclease-free water. Real-time quantitative RT-PCRs were performed on cDNA samples using gene-specific primers (Supplementary Table 1) as described in Campoli et al. (2012a) and Bi et al. (2019). Two technical replicates were used for each sample and non-template controls were included. Starting concentration of the target transcripts were calculated according to the absolute quantification method with primer efficiency correction based on the titration curve for each target gene using the LightCycler 480 Software (Roche; version 1.5) and normalized against the geometric mean of the reference genes *HvActin, HvGAPDH* and *HvADP* (Supplementary Table 1).

### Multiple protein alignment

Amino acid sequences from selected FT homologs from Arabidopsis, onion (*Allium cepa)*, sugar beet (*Beta vulgaris*), longan (*Dimocarpus longan*), soybean (*Glycine max*), sunflower (*Helianthus annuus*), tobacco (*Nicotiana tabacum*), sugarcane (*Saccharum* spp.), and Norway spruce (*Picea abies*) were selected, which have been extensively studied in Wickland and Hanzawa (2015). This selection was expanded with FT homologs from rice (*Oryza sativa*) and wheat (*Triticum aestivum*), of which FT function has been characterized, and all barley FT homologs (Halliwell et al., 2016). Multiple protein alignments of full coding regions of PEBP containing FT-like proteins were created using ClustalW with default settings on the MUSCLE homepage (Multiple Sequence Comparison by Log-Expectation, https://www.ebi.ac.uk/Tools/msa/muscle/, 21.11.2017). The protein sequences were sorted into seven groups according to function and/or species. FT and TFL1 proteins from Arabidopsis were considered as group 1 and used as a reference. FT-like proteins from other cereal species with close homology to barley FT-like proteins, as determined in a phylogenetic analysis by Halliwell et al. (2016), were collected in group 2. Group 2 FT-like proteins are described as floral promoters and were included to confirm the conservation of the amino acid pattern, which is critical to determine FT-like function in cereal monocots. Furthermore, barley FT-like proteins with described inductive (Group 3) or repressive function on flowering time (Group 4), as well as FT-like proteins from various species with described function as floral promoters (Group 6) or floral repressors (Group 7) (Wickland and Hanzawa, 2015) were arranged accordingly in the alignment. All remaining barley FT-like genes with unknown function were included in group 5. Corresponding protein IDs or gene model information are listed in Supplementary Table 2.

### Statistical analysis

Data was visualized and analysed using the RStudio software (version 1.0.153, R Development Core Team, 2008). Significant differences in flowering time, developmental stage of the SAM, morphological phenotypes and gene expression levels between each of the *Ubi::HvFT4* families, the null control and the wild-type Golden Promise were identified by one-way analysis of variance (one-way ANOVA) followed by Tukey’s multiple comparison test (Tukey HSD) with p-value adjustment. The statistical significance of differences in *HvFT4* expression levels between the two phenotypic categories of *Ubi::HvFT4* (Category I and 0) was determined using a Welch Two Sample t-test. The significance level was α = 0.05 for all tests.

## Results

### Overexpression of *HvFT4* prolongs the vegetative phase and delays flowering

The constitutive overexpression of *HvFT4* in the background of Golden Promise significantly delayed time to flowering (Figure 1A). The three independent transgenic *Ubi::HvFT4* families flowered on average 71 days after emergence (DAE), while Golden Promise and the null segregants required on average only 59 days to flowering. Furthermore, 35 % of transgenic *Ubi::HvFT4* plants exhibited an impaired main shoot development (Figure 1B, D). These plants, herein referred to as category 0, remained small and the main shoot apex was aborted during late stem elongation and failed to flower, while the remaining 65 % of the transgenic lines (category 1) eventually flowered and formed a spike (Figure 1B, D). *HvFT4* was strongly upregulated in the transgenic lines compared to the control lines (Figure 1C). The expression of *HvFT4* was 2-fold higher in the leaves of category 0 plants (data not shown) compared to the expression levels of *HvFT4* in plants of category I (p = 5.16×10-06, Welch Two Sample t-test). Thus, upregulation of *HvFT4* expression led to a delay in flowering or even the abortion of the main shoot.

**Figure 1:**
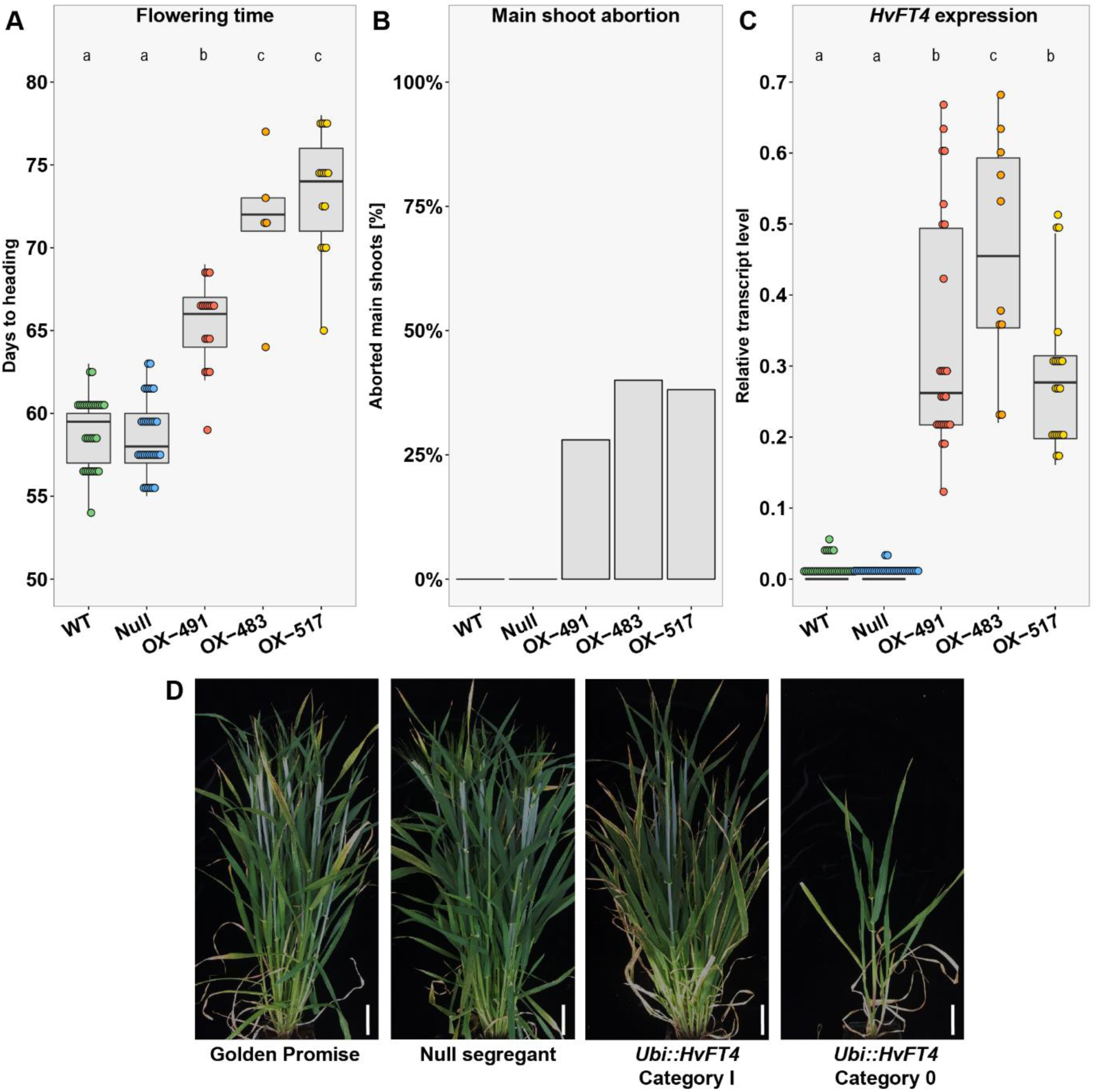
Overexpression of *HvFT4* delays flowering and leads to premature main shoot abortion. **A:** Flowering time was measured in days from emergence until heading of the main shoot (Zadoks stage 49) (Zadoks et al., 1974). **B:** Proportion of prematurely aborted main shoots [%]. **C:** *HvFT4* expression was measured in the youngest fully emerged leaf of the main shoot 12 DAE at zeitgeber time T10 under LDs (16h light/8h night). The expression level of each gene was normalized against the geometric mean of the reference genes *HvActin, HvGAPDH* and *HvADP.* Each dot represents the values obtained from a single plant. Statistical differences (p ≤ 0.05) between families were calculated by one-way analysis of variance (one-way ANOVA) followed by Tukey’s multiple comparison test (Tukey HSD). **D:** Representative plants of every genotype. *UBI::HvFT4* plants showed two distinct phenotypes (Category I and Category 0). Category 0 plants showed impaired development and higher levels of *HvFT4* expression compared to Category I plants. Scale bar represents 5 cm. **WT** = Golden Promise, **Null** = null segregant, **OX-491** = *Ubi::HvFT4*-491, **OX-483** = *Ubi::HvFT4*-483, **OX-517** = *Ubi::HvFT4*-517.

We then examined the effects of *HvFT4* overexpression on individual phases of pre-anthesis development. For this purpose, developing primary shoots of one transgenic family (OX-517) and both control lines (WT and null) were dissected and inflorescence development was evaluated according to the Waddington scale (Waddington et al., 1983). This scale describes the development of the inflorescence and the most advanced floret primordium and carpel within the inflorescence. The development of the first spikelet primordia on the shoot apex at the double ridge stage (W1.5-W2.0) marks the transitions to a reproductive shoot apical meristem (SAM). The first floral organ primordia differentiate at the stamen primordium stage (W3.5), when also stem elongation initiates. Anthesis and pollination of the most advanced floret take place at the Waddington stage W10.0.

Apical meristems of the control plants had already initiated spikelet primordia (W2.0) 21 DAE, whereas the transgenic plants required eight days more to reach the same stage (Figure 2). Stem elongation (W3.5) was initiated in the transgenic lines 42 DAE, whereas the control lines had reached the same stage seven days earlier. Golden Promise lines flowered 64 DAE, while the transgenic line flowered only 71 DAE (Figure 2A). Consequently, the delay in flowering time was primarily due to a prolonged vegetative phase in the *Ubi::HvFT4* plants (Figure 2B).

**Figure 2:**
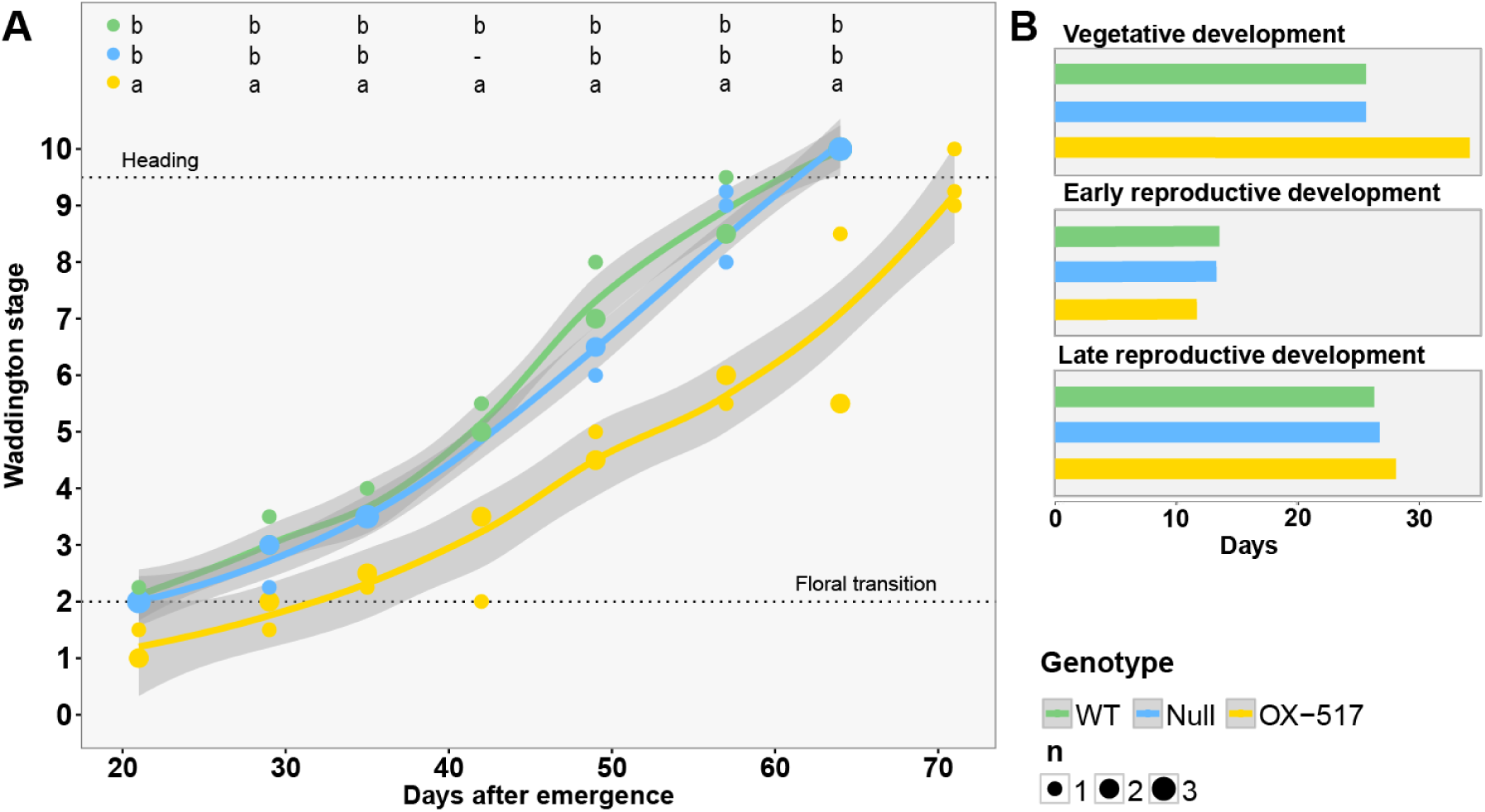
Overexpression of *HvFT4* delays development of the main shoot apex. **A:** Microscopic development of the MSA was scored every seven days according to the Waddington scale (Waddington et al., 1983). Three plants per genotype were dissected at each time point. Polynomial regression models at 95 % confidence interval (Loess smooth line) are shown. Statistical differences (p ≤ 0.05) at each time point were calculated by one-way analysis of variance (one-way ANOVA) followed by Tukey’s multiple comparison test (Tukey HSD). **B:** Duration of three distinct developmental phases were extracted graphically from figure 9 A. The length of the vegetative phase was measured as the days from germination until Waddington stage 2.0, the early reproductive phase was accounted as the time from Waddington stage 2.0 until 3.5 and the time from Waddington stage 3.5 until heading was considered as the late reproductive phase. **WT** = Golden Promise, **Null** = null segregant, **OX-517** = *Ubi::HvFT4*-517.

The delay in reproductive development in Ubi::*HvFT4* was correlated with a reduced harvest index (HI) determined as the weight of the filled spikes in proportion to the total shoot biomass of the plant (Figure 3A). Null segregants and Golden Promise had a harvest index of on average 0.34, whereas *Ubi::HvFT4* plants had a harvest index of 0.17 (OX-491), 0.05 (OX-483), and 0.1 (OX-517), which corresponds to a reduction in HI of 49 % to 84 % (Figure 3A). This reduction in HI was mainly caused by a decrease in reproductive biomass. The number of florets and the number of grains on the main spike were significantly reduced in transgenic plants compared to the control plants (Figure 3B, Supplementary Figure 1C). The reduction in overall spike fertility was mainly attributed to reduced grain set in the central zone of the spike in *Ubi::HvFT4* plants, whereas the apical and basal part was also partly sterile in control plants (Supplementary Figure 2). Interestingly, *Ubi::HvFT4* plants developed significantly fewer tillers at flowering compared to control plants even though they initiated spikelet primordia significantly later and consequently formed more leaves on the main culm than the wild-types (Figure 3C, Supplementary Figure 1D). Additionally, the proportion of tillers with spikes containing at least one grain was significantly lower in *Ubi::HvFT4* compared to control plants (Figure 3D). However, *HvFT4* overexpression had no effects on plant height, peduncle extrusion or leaf size in this study (Supplementary Figures 1A, B and 3). Finally, thousand grain weight (TGW), grain width and grain size were reduced in *Ubi::HvFT4* compared to the controls (Supplementary Figure 4). Taken together, overexpression of *HvFT4* decreased reproductive biomass by reducing tiller and spike number, grain number per spike and grain size.

**Figure 3:**
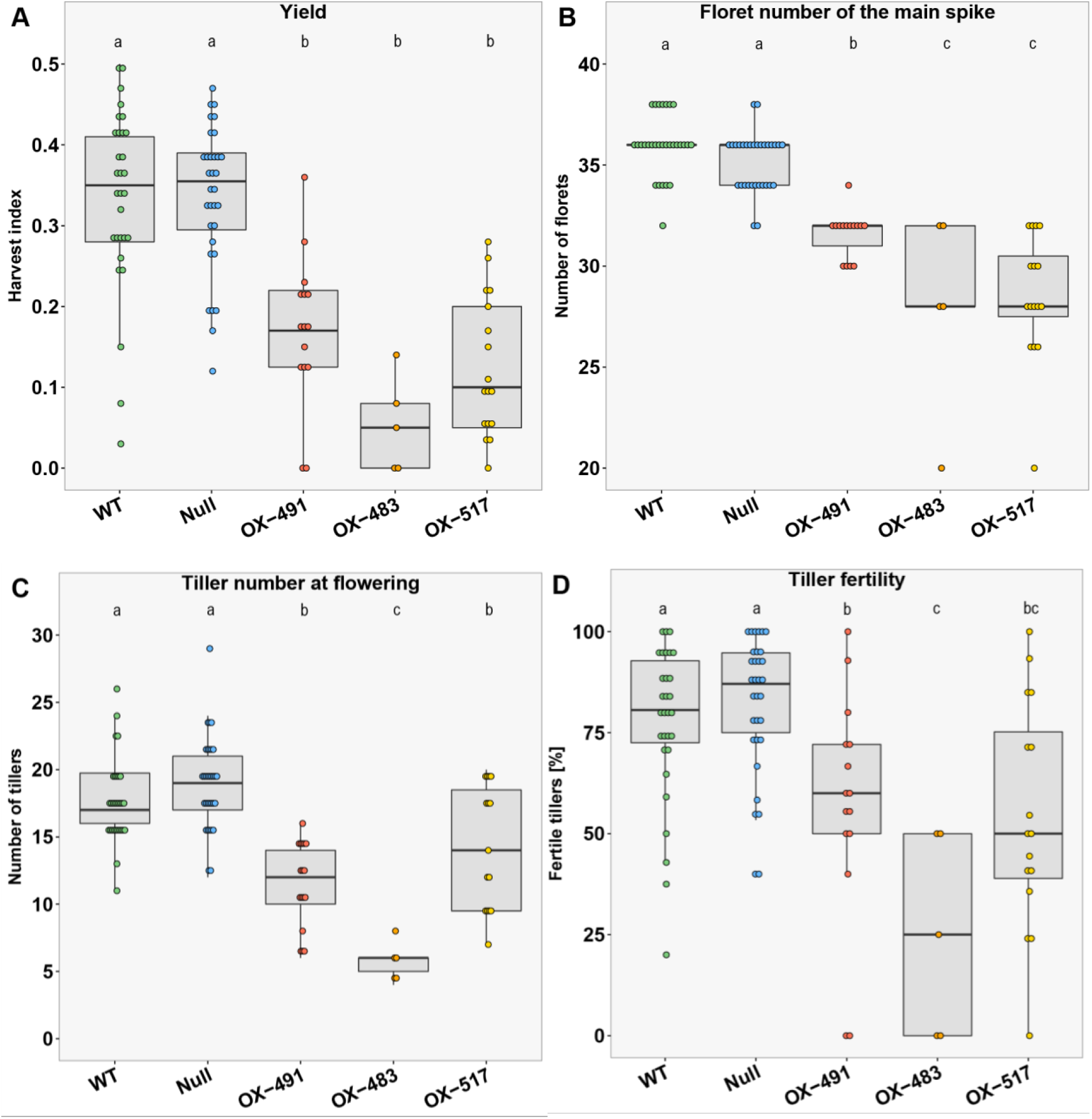
Overexpression of *HvFT4* reduces yield, tiller number and fertility. **A:** Harvest index 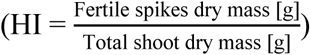, **B**: Total number of florets on the main spike, **C:** number of tillers and **D:** fertile tillers [%] (= spike with at least one grain) were recorded at plant maturity. Each dot represents the values obtained from a single plant. Statistical differences (p ≤ 0.05) between genotypes were calculated by one-way analysis of variance (one-way ANOVA) followed by Tukey’s multiple comparison test (Tukey HSD). **WT** = Golden Promise, **Null** = null segregant, **OX-491** = *Ubi::HvFT4*-491, **OX-483** = *Ubi::HvFT4*-483, **OX-517** = *Ubi::HvFT4*-517.

### Overexpression of *HvFT4* reduced expression of the *AP1*-like flowering promoters in the leaves and the shoot apical meristem

To understand the role of HvFT4 in the control of flowering and identify genes which are regulated by HvFT4, the influence of *HvFT4* overexpression on the expression levels of known flowering time regulators was analysed in the leaves and developing inflorescences.

Native *HvFT4* expression levels were very low in the leaf and strongly upregulated in the transgenic plants (Figure 4A, 5A). *HvFT1* levels in the leaves were low in all plants and only started to increase after 49 DAE in the WT plants and after 60 DAE in the transgenic plants (Figure 4B). Similarly, *HvFT2* expression levels in the leaves were below the detection limit until 42 DAE and 57 DAE when levels started increasing in the WT and transgenic lines, respectively (Figure 4C). Expression levels of *HvFT1* and *HvFT2* were significantly different between *Ubi::HvFT4* and control plants at late reproductive stages (Figure 4B, C). Overexpression of *HvFT4* had no significant effects on *HvFT3* and *Ppd-H1* expression levels in the leaves (Figure 4D, E). We further analysed the expression of the *AP1*-like genes *VRN-H1, HvBM3* and *HvBM8* which are putative targets of *HvFT1* and *HvFT3* and promote floral development of barley (Hemming et al., 2008; Digel et al., 2015; Mulki et al.; 2018). *VRN-H1* was significantly downregulated in the leaves of transgenic plants at all time points (Figure 4F). *HvBM3* transcript levels were low, but detectable in the leaves of all genotypes until 35DAE and 50 DAE when expression levels increased in the WT and transgenic plants, respectively. *HvBM3* expression levels were significantly lower in transgenic plants compared to control plants at most time points (Figure 4G). *HvBM8* expression levels in the leaves were below the detection levels until 42 DAE when they were strongly upregulated in the WT, but not in transgenic plants, where *HvBM8* expression levels remained low until 60 DAE (Figure 4H). In the inflorescence, *HvFT1* and *HvFT2* levels were below the detection limit at W3.5-4.5, but detectable at W6.0-7.0 when expression levels were higher in WT than transgenic plants (Figure 5B, C). Expression levels of *VRN-H1* were significantly downregulated in the transgenic versus WT plants at W6.0-7.0, while *HvBM3* and *HvBM8* expression levels were significantly reduced in transgenic compared to WT plants at W3.5-4.0 and W6.0-7.0 (Figure 5D-F). Additionally, the expression level of the barley MADS-box transcription factors and putative floral repressors *HvBM1* and *BM10* was measured (Hartmann et al., 2000; Trevaskis et al., 2007). While expression of *HvBM1* and *BM10* was significantly upregulated in the leaf at late developmental stages, no significant differences were detected in the inflorescence (Figure 4I, J, Figure 5G). *Ppd-H1* was expressed in the leaf and inflorescence, but no significant differences in expression levels were observed (Figure 5H). As overexpression of *HvFT4* caused a strong reduction in tillering, we also assayed expression of *INTERMEDIUM-C (INT-C)*, which is a repressor of axillary bud outgrowth (Ramsay et al., 2011; Liller et al., 2015). However, no effect of *HvFT4* overexpression on the expression of *INT-C* was observed neither in the leaf nor inflorescence (Figure 5 I).

**Figure 4:**
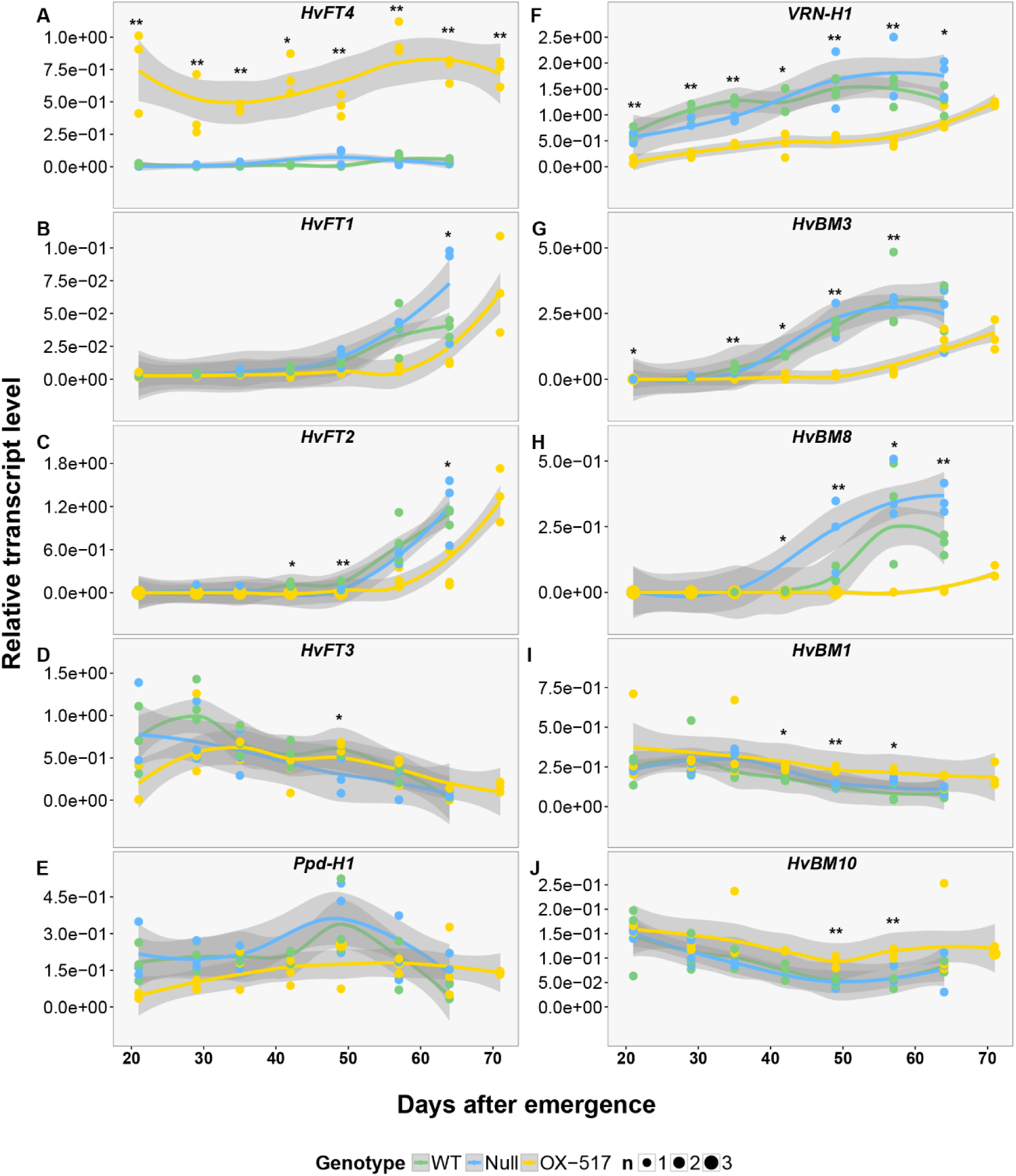
Temporal expression pattern of flowering time genes in the leaves of *Ubi::HvFT4*, null segregant and Golden Promise. Temporal expression of flowering time genes was assayed in the youngest fully emerged leaf of the main shoot of three plants of every genotype, every seven days at zeitgeber time T10 under longday conditions (16h light/8h night). The expression level of each gene was normalized against the geometric mean of the reference genes *HvActin, HvGAPDH* and *HvADP*. Polynomial regression models at 95 % confidence interval (Loess smooth line) are shown. For better visualization asterisks instead of letters were used to indicate significant differences (**\***: OX-517 differed significantly from one control line (WT **or** null), **\*\***: OX-517 differed significantly from both control lines (WT **and** null). Corresponding letters are given in Supplementary Table 2. Statistical differences (p ≤ 0.05) at each time point were calculated by one-way analysis of variance (one-way ANOVA) followed by Tukey’s multiple comparison test (Tukey HSD). (**WT** = Golden Promise, **Null** = null segregant, **OX-517** = *Ubi::HvFT4*-517).

**Figure 5:**
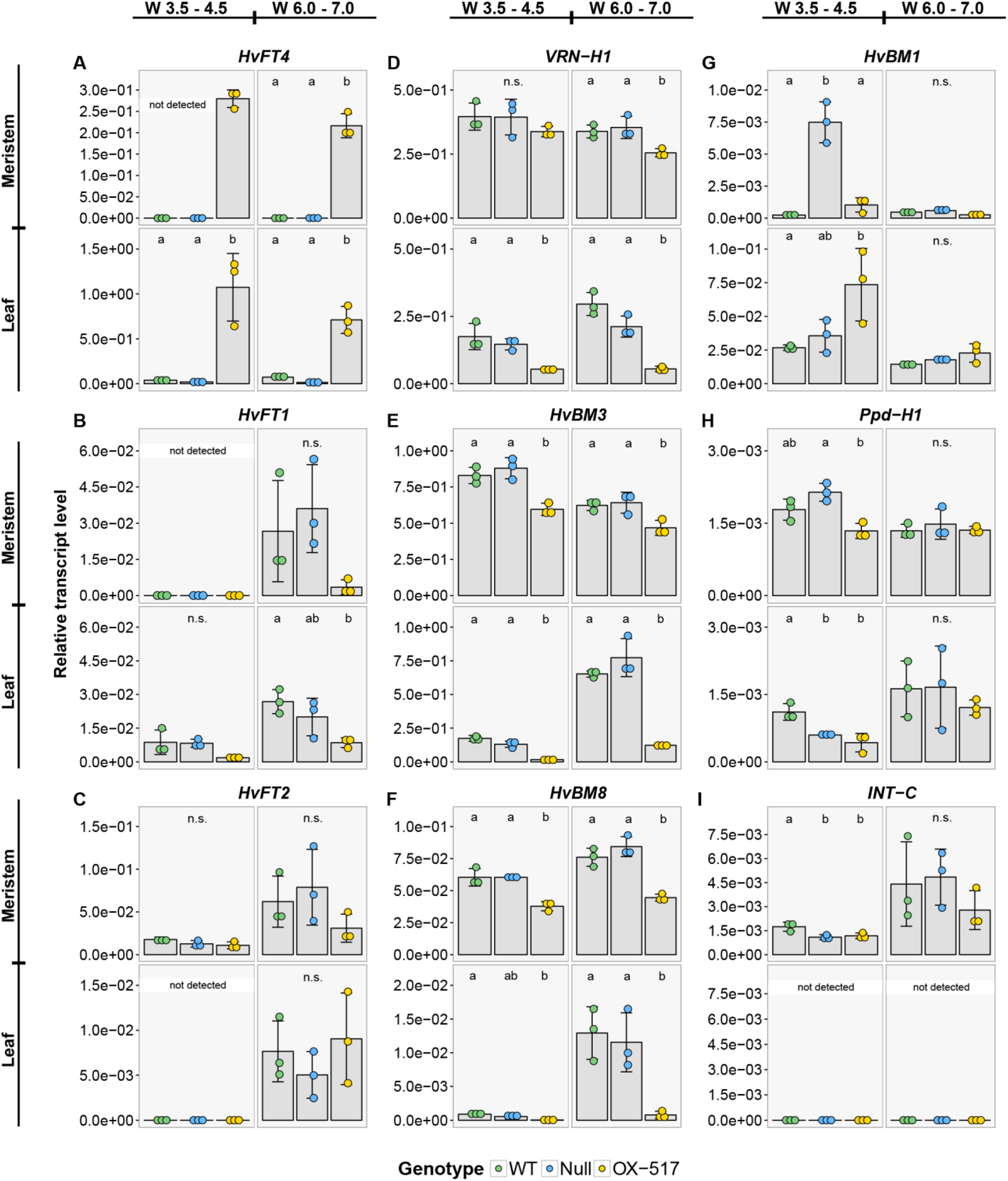
Influence of *Ubi::HvFT4* on the expression level of flowering-related genes in the leaves and meristem at Waddington stages 3.5-4.5 and 6.0-7.0. Expression of flowering time genes was assayed at two stages during reproductive development (W3.5-W4.5 and W6.0-W7.0) at zeitgeber time T10 under long day conditions (16h light/8h night). The expression level of each gene was normalized against the geometric mean of the reference genes *HvActin, HvGAPDH* and *HvADP*. Each dot represents a pool of main shoot apices or leaf material from five single plants. Bars represent mean ± SD. Statistical differences (p ≤ 0.05) between genotypes where determined by one-way analysis of variance (one-way ANOVA) followed by Tukey’s multiple comparison test (Tukey HSD). (**WT** = Golden Promise, **Null** = null segregant, **OX-517** = *Ubi::HvFT4*-517).

Taken together, overexpression of *HvFT4* was associated with a downregulation of *HvFT1* and *HvFT2* in the leaf, of *VRN-H1, HvBM3* and *HvBM8* in the leaf and inflorescence, and an upregulation of *HvBM1* and *BM10* in the leaf.

### Multiple sequence alignment reveals amino acid substitutions in *HvFT4* in conserved motifs

The phenotypic and expression analysis suggested that HvFT4 acts as a repressor of flowering through downregulating of floral promotors in the leaf and inflorescence. It is well known that small changes in individual amino acid residues of *FT*-like genes determine whether they act as repressors or activators of flowering (Ahn et al., 2006; Kaneko-Suzuki et al., 2018). We aimed to identify amino acid residues which might explain the repressive function of HvFT4 by aligning the HvFT4 protein sequence with protein sequences of *FT*-like genes from various species (Supplementary Table 3).

Sequence comparison revealed that the amino acid residues R63, T67, P94, F101 and R130, previously determined to be critical for FT-14-3-3 interaction in wheat and rice (Taoka et al., 2011; Li et al., 2015), are conserved among barley HvFT1 to HvFT4 (Figure 6). In Arabidopsis, the signatures Y85 and Q140 differentiate the floral promotor FT from its sister protein TFL1 which acts as a repressor of flowering (Hanzawa et al., 2005; Ahn et al., 2006). However, HvFT4 shares these functionally important FT signatures with FT and floral repressors from other species, but not the repressor TFL1 (Figure 6). The LYN triad, which is immediately adjacent to and makes contact with the segment B external loop in Arabidopsis (Ahn et al., 2006), is conserved in HvFT1, HvFT2 and other FT-like proteins which function as floral promoters (Figure 6). It is, however, more variable in proteins characterised as floral repressors including HvFT4 (Figure 6). Remarkably, HvFT4 shows the same amino acid pattern (L159R, Y150R, N150) as ScFT1, another putative FT-like floral repressor from the Poaceae species sugarcane (*Saccharum* spp.) (Coelho et al., 2014).

**Figure 6:**
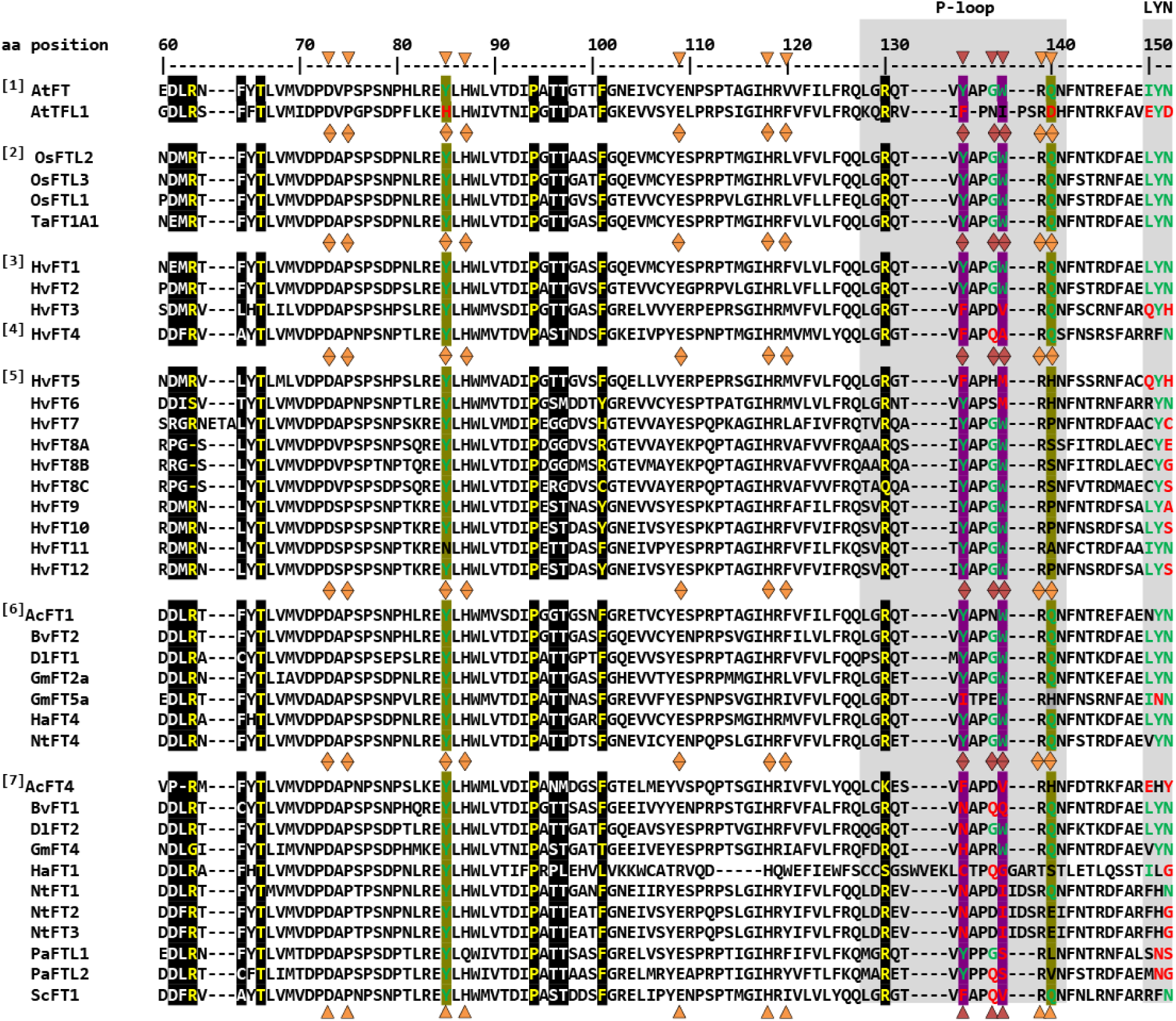
Inter- and intra-species comparison of amino acid sequences of PEBP homologues. Group **[1]**: FT and TFL1 protein from Arabidopsis, group **[2]**: Inductive rice and wheat FT-like proteins, group **[3]**: Inductive barley FT-like proteins, group **[4]**: Repressive barley FT-like proteins, group **[5]**: Uncharacterised barley FT-like proteins, group **[6]**: Inductive FT-like proteins from other species, group **[7]**: Repressive FT-like proteins from other species. The **P-loop** (position 128 – 141) and **LYN triad**, (position 150 – 153) of exon 4 are boxed in grey. The amino acid residues at position 85 and 140 unambiguously distinguish between FT- and TFL1 homologues are highlighted in olive (Hanzawa et al., 2005; Ahn et al., 2006). The amino acid residues at position 134 and 138, which are associated with inductive versus repressive function of FT-like proteins are highlighted in purple (Wickland and Hanzawa, 2015). The amino acid residues at position 134, 138 and 137, which are responsible for the opposing functions of BvFT2 and BvFT1 in beet, are marked with a red triangle (Pin et al., 2010). Amino acid residues, which are specifically attributed to FT-like genes **and/or** are associated with flowering promotion, are displayed in green letters. Amino acid residues, which are specifically attributed to TFL1-like genes **and/or** are associated with flowering repression, are displayed in red letters (Pin et al., 2010; Klintenäs et al., 2012; Wickland and Hanzawa, 2015). Amino acid residues with no characterised effect on flowering time are depicted in black letters. Amino acid residues lining the ligand-binding site as shown by Ahn et al. (2006) marked with an orange triangle. Amino acid residues located at the binding interface with 14-3-3 protein are highlighted in black with white characters. Critical amino acid residues, which abolished protein interactions between 14-3-3 proteins and TaFT1 or OsFTL2 respectively in yeast two-hybrid assays when mutated are shown in yellow letters (Taoka et al., 2011; Li et al., 2015). All positions given refer to the amino acid position in Arabidopsis FT protein. Complete protein sequences were aligned with ClustalW. For better visualization only the part of the PEBP domain is shown, which includes amino acid residues critical for PEBP protein function.

Taken together, the sequence comparison suggested candidate amino acid residues that might confer the repressive activity of HvFT4. Which of these residues is causative still needs to be elucidated.

## Discussion

In this study, *HvFT4* was overexpressed in the genetic background of the spring cultivar Golden Promise under LDs to investigate the influence of HvFT4 on flowering time and inflorescence development in barley. Overexpression of *HvFT4* prolonged vegetative growth and delayed spikelet initiation. The effects of HvFT4 were antagonistic to those of HvFT1, HvFT2 and HvFT3, which promote spikelet initiation and inflorescence development in barley (Digel et al., 2015, Mulki et al., 2018; Shaw et al. 2019). Typically, a delay in reproductive development and reduction in apical dominance are associated with an increase in the number of spikelets and tillers in barley (Digel et al., 2015; Bi et al., 2019). However, overexpression of *HvFT4* delayed reproductive development, decreased the number of spikelets on the main culm and reduced the number of tillers and grain weight. Consequently, overexpression of *HvFT4* had negative pleiotropic effects on a number of reproductive traits in barley. These effects of HvFT4 on apical, axillary and spikelet meristems suggested that HvFT4 plays a role in the development of different shoot meristems. Tsuji et al. (2015) already showed that *Hd3a*, an *HvFT1* orthologue in rice, not only promotes the development of the apical meristem, but also that of lateral buds and therefore influences branching in rice. Similarly, mutations in *CENTRORADIALIS* (*HvCEN*), a barley homolog of the floral repressor *TFL1*, reduced tiller number, spikelet and grain number per spike (Bi et al., 2019). Furthermore, a key regulator of axillary bud outgrowth is *TEOSINTE BRANCHED 1 (TB1)*. Homologs of the maize (Zea mays) *TB1* gene, *INTERMEDIUM-C* (*INT-C*) in barley and *BRANCHED 1 (BRC1)* Arabidopsis, suppress axillary bud outgrowth (Doebley et al., 1997; Hubbard et al., 2002). In Arabidopsis, BRC1 was shown to interact with the FT protein to delay floral transition in the axillary meristems (Niwa et al., 2013). Similarly, it was recently shown that in wheat the TB1 protein interacts with FT1 and that increased dosage of TB1 alters inflorescence architecture and tiller number (Dixon et al., 2018). While overexpression of *HvFT4* did not affect *INT-C* expression, HvFT4 protein might interact with INT-C and thereby affect the TB1/FT1 dosage und thus tiller outgrowth and inflorescence architecture. Similarly, it was shown in Arabidopsis that overexpression of *BROTHER OF FT AND TFL1 (BFT)*, which shares a higher sequence similarity with FT than with TFL1, resulted in late flowering and suppression of axillary inflorescence growth (Kobayashi et al., 1999; Yoo et al., 2010). However, the results obtained by the constitutive overexpression of *HvFT4* need to be interpreted with caution. It is also possible that the reduced number of tillers observed in *Ubi::HvFT4* plants resulted from ectopic expression of *HvFT4* in tissue where it is usually not expressed. For example, we did only observe expression of the native *HvFT4* in the leaves, but not in the inflorescence.

Overexpression of *HvFT4* was associated with a strong downregulation of the *AP1-*like genes *VRN-H1, HvBM3* and *HvBM8* in the leaves and meristem. In rice, simultaneous knockdown of *OsMADS14* (*VRN1, FUL1*), *OsMADS15* (*BM3, FUL2*) and *OsMAD18* (*BM8, FUL3*) impaired spikelet development and resulted in floral reversion (Kobayashi *et al.*, 2012). Similarly, triple wheat *vrn1ful2ful3* mutants developed vegetative tillers instead of spikelets (Li *et al.*, 2019). Low expression levels of *VRN-H1, BM3* and *BM8*in *Ubi::HvFT4* plants might have caused the delay in spikelet initiation However, and reduced floret fertility and grain set. It has already been demonstrated that *FT1, FT2*, and the *BM* genes affect floret fertility and grain set (Digel et al., 2015; Ejaz et al., 2017; Shaw et al., 2019). We therefore conclude that the downregulation of *BM*-like genes and consequently *HvFT1* and *HvFT2* in the inflorescence has contributed to the reduction in fertility and grain set of the main and axillary shoots.

Evidence from Arabidopsis shows that FT and TFL1 act as antagonists by competing for the same binding partners (Ahn et al., 2006). It was proposed that FT recruits a transcriptional activator, whereas TFL1 recruits a transcriptional repressor (Hanzawa et al., 2005; Ahn et al., 2006; Hanano and Goto, 2011; Ho and Weigel, 2014). HvFT4 might also compete with HvFT1, HvFT2 and HvFT3 for binding partners and thus control the expression of putative target genes such as the *BM* genes. Studies in maize and Brachypodium further support this hypothesis. Maize plants with reduced expression of the floral promoter *FT*-like gene *Zea CENTRORADIALIS 8* (*ZCN8*) resembled the phenotype of plants which overexpressed the floral repressor *TFL1*-like Zea *CENTRORADIALIS 2* (*ZCN2*), suggesting that both proteins compete for the same interaction partners (Danilevskaya et al., 2011). In Brachypodium an SD-induced FLOWERING LOCUS T ortholog, FT-like 9 (FTL9), promotes flowering in SDs but inhibits flowering in LDs (Qin et al., 2019). Both proteins could interact with FD1 to form a flowering activation complex (FAC) but with lower activity of FTL9-FAC than of FT1-FAC. This likely resulted in a positive role for FTL9 in promoting floral transition under SDs when FT1 is not expressed, but a dominant-negative role when FT1 accumulates under LDs. Consequently, we propose that HvFT4 may also compete with *HvFT*-like proteins that act as floral activators to repress flowering under LDs. However, further experimental evidence is needed to analyse the activity and binding partners of the HvFT4 protein (Li et al., 2015).

Sequence comparison of FT-like proteins suggested several residues that might be causal for the repressive function of HvFT4. The amino acid residues critical for FT-14-3-3 interaction were conserved in barley HvFT4, suggesting that HvFT4 possesses 14-3-3 binding capacities and engages into a normal florigen activation complex (FAC) formation. Indeed, HvFT1, HvFT3, and HvFT4 were shown to interact with the same 14-3-3 proteins in a yeast-two-hybrid assay (Li et al., 2015). Secondly, HvFT4 and HvFT3 proteins carry amino acid substitutions within their P-loop among others at the amino acid positions 134, 137, and 138. Interestingly, these three amino acids were identified to be the major cause of antagonistic functions of two FT-like proteins in beet (Pin et al., 2010). The exchange of the external loop of BvFT1 for the P-loop of BvFT2, with the three altered amino acids, converted BvFT1 from a floral repressor to an activator (Pin et al., 2010). Remarkably, HvFT4, but not HvFT3, shares a glutamine at position 137 with the flowering repressor BvFT1. However, additional experiments with single amino acid swapping indicated that Y134, W138 and Y85 are essential for the inductive function of FT-like genes in beet (Klintenäs et al., 2012).

Furthermore, HvFT3 and HvFT4 do not carry the conserved amino acids Y134 and W138, which are associated with FT-like flowering inductive function, but hydrophobic amino acids they share with the flowering repressor TFL1 (Wickland and Hanzawa, 2015). It is still not known which amino acid position is critical for flowering promotive or repressive function in barley and which substitutions are tolerated or lead to modification of the protein function. However, HvFT4 sequence features correspond to those of other known repressor *FT* genes and are indicative of candidate amino acid residues that might confer the repressive activity of HvFT4.

Taken together, our study provided phenotypic and molecular data that indicates *HvFT4* may act as a repressor of reproductive development in barley. Plants overexpressing *HvFT4* displayed a delay in reproductive development, a reduction in spikelet and tiller number and floret fertility, and a downregulation of genes promoting spikelet and floral development.

## Supplemental Data

Supplementary Table 1: List of primers used in this study.

Supplementary Table 2: Protein IDs or gene model information used for the multiple sequence alignment.

Supplementary Table 3: Significant differences corresponding of the temporal and developmental expression pattern of flowering time genes in the leaves of *Ubi*::*HvFT4*, null segregant and Golden Promise (Figure 4).

Supplementary Figure 1: Overexpression of *HvFT4* decreases tillering and increases the number of leaves on the main shoot, but does not influence plant height and peduncle extrusion.

Supplementary Figure 2: Overexpression of *HvFT4* reduces floret fertility of the main shoot spike.

Supplementary Figure 3: Overexpression of *HvFT4* does not influence leaf size. Supplementary Figure 4: Overexpression of *HvFT4* reduces grain weight and width.

## Conflict of Interest

The authors declare that the research was conducted in the absence of any commercial or financial relationships that could be construed as a potential conflict of interest.

## Author Contributions

R.P., F.T. and M.v.K. planned and designed the research, R.P. and F.T. performed the experiments and analyzed the data, R.P., F.T. and M.v.K. wrote and revised the manuscript.

## Acknowledgments

We cordially thank Kerstin Luxa, Caren Dawidson and Andrea Lossow for excellent technical assistance. Agatha Walla is acknowledged for critically reading this manuscript. This work was funded by the Max Planck Society and the Deutsche Forschungsgemeinschaft (DFG, German Research Foundation) under Germany’s Excellence Strategy – EXC-2048/1 – Project ID: 390686111.

